# Gene interaction enrichment analysis for transcriptomic data with GREA

**DOI:** 10.1101/2025.04.15.649030

**Authors:** Xiaoyu Liu, Anna Jiang, Chengshang Lyu, Lingxi Chen

## Abstract

Gene Set Enrichment Analysis (GSEA) is a cornerstone for interpreting gene expression data, yet traditional approaches overlook gene interactions by focusing solely on individual genes, limiting their ability to detect subtle or complex pathway signals. To overcome this, we present GREA (Gene Interaction Enrichment Analysis), a novel framework incorporating gene interaction data into enrichment analysis. GREA replaces the binary gene hit indicator with an interaction overlap ratio, capturing the degree of overlap between gene sets and gene interactions to enhance sensitivity and biological interpretability. It supports three enrichment metrics: Enrichment Score (ES), Enrichment Score Difference (ESD) from a Kolmogorov-Smirnov-based statistic, and Area Under the Curve (AUC) from a recovery curve. GREA evaluates statistical significance using both permutation testing and gamma distribution modeling. Benchmarking on transcriptomic datasets related to respiratory viral infections shows that GREA consistently outperforms existing tools such as blitzGSEA and GSEApy, identifying more relevant pathways with greater stability and reproducibility. By integrating gene interactions into pathway analysis, GREA offers a powerful and flexible tool for uncovering biologically meaningful insights in complex datasets. The source code is available at https://github.com/compbioclub/GREA.

## Introduction

Gene set enrichment analysis (GSEA) has long been central to interpreting transcriptomic data by elevating inference from individual genes to pathways and biological processes (1). Building on the classical GSEA framework (2), which ranks genes by differential expression and uses a Kolmogorov–Smirnov-like (KS-like) (3) running-sum statistic to test whether set members accumulate at the extremes of a ranked list, numerous variants have expanded both scope and efficiency. blitzGSEA (4) accelerates significance estimation by fitting the null distribution with a Gamma model rather than relying solely on extensive permutations. Single-sample extensions, including ssGSEA (5) and GSVA (6), enable pathway activity profiling at the individual—sample level without predefined case–control labels: ssGSEA computes sample-wise enrichment from absolute expression, while GSVA models gene expression distributions nonparametrically and applies KS-style scoring and enrichmentscore differences. Practical Python implementations such as GSEApy (7) further democratize these approaches across pathway libraries and statistical settings.

Despite these advances, most enrichment methods (Table 1) implicitly assume gene independence, emphasizing marginal changes in single genes while underrepresenting the coordinated interactions that drive cellular function. Biological processes are emergent, arising from networks of gene–gene relationships. Ignoring this structure can limit sensitivity to subtle but coordinated perturbations and may hinder interpretability, especially when regulatory effects are distributed across interacting partners. A persistent gap remains: incorporating interaction information directly into the enrichment machinery in a way that is both statistically robust and computationally tractable.

**Table 1.** Functionality comparison of enrichment methods. DT: Differential Test, WKS: Weighted Kolmogorov-Smirnov, RC: Recover Curve, ES: Enrichment Score, NES: Normalized Enrichment Score, ESD: Enrichment Score Difference, AUC: Area under Curve, L: Low, M: Moderate, H: High, A: Accurate.

|  |  | GREA | GSEApY | blitzGSEA |
| --- | --- | --- | --- | --- |
| Rank Signature | Phenotype-specific | ✓ | ✓ | ✓ |
|  | Rank unit | gene interaction | gene | gene |
| Enrichment | Rank value | DT log2FC/z-score of entropy score | DT log2FC/z-score of gene expression | DT log2FC/z-score of gene expression |
|  | WKS approach | ES, ESD | ES, ESD | ES, ESD |
|  | RC approach | AUC | - | - |
| Significant Testing | Permutation | ✓ | ✓ | - |
|  | Analytical | Gamma | - | Gamma |
| Software | Efficiency | H,A | M | H,A |
|  | Access | Python | Python | Python |

To fill this gap, we introduce GREA (Gene Interaction Enrichment Analysis), a general framework that performs enrichment over ranked gene–gene interactions rather than isolated genes. Conceptually, GREA replaces the binary “gene hit” indicator with an interaction overlap ratio that quantifies the degree of alignment between a target gene set and each interaction. This redefinition enables the application of KS-based running-sum statistics to interaction-ranked lists while preserving the familiar intuition of early-versus-late accumulation. To capture early recovery of pathway-relevant interactions, GREA also implements a recovery-curve formulation with area under the curve (AUC) scoring, which can increase sensitivity when informative interactions cluster near the top of the ranking. For significance assessment, GREA combines permutation-based testing with Gamma-model fitting of the null distribution to stabilize p-values and reduce variance, particularly in settings with limited samples, sparse signals, or heterogeneous preprocessing pipelines where pure permutation can be noisy or computationally expensive. This hybrid strategy yields better-calibrated and more reproducible inference across diverse analytic contexts.

We systematically benchmarked GREA on public transcriptomic datasets spanning respiratory viral infection and multiple cancer types. Across preprocessing strategies, scoring functions, and statistical paradigms, GREA consistently delivered lower *p*-value variance than established benchmarks and robustly recovered known pathways reported in the literature. In respiratory infection datasets, GREA recapitulated expected immune and antiviral processes, demonstrating concordance with prior findings. In cancer analyses, GREA identified canonical oncogenic pathways and, importantly, revealed strong enrichment signals specific to gene–gene interactions that were attenuated or missed by single-gene methods—highlighting the added value of modeling coordinated biology. Together, these results position GREA as a practical, interaction-aware enrichment framework that enhances stability, sensitivity, and interpretability in pathway analysis.

## Methods

### Algorithm of GREA

#### Problem Formulation

Consider a gene regulatory network (GRN) represented as a graph *G* = (𝒱, *ℰ*), where vertices *v* ∈ *𝒱* correspond to genes and edges (*u, v*) ∈ *ℰ* represent gene interaction between *u* and *v*. There are *N* = |*ℰ*| gene interactions, and each interaction *ℐ* = (*u, v*) is associated with a quantitative rank score *x*_*u,v*_. When phenotypic labels are available, the score is phenotype-specific, derived from gene expression data conditioned on the phenotype. Otherwise, the score is specific to each sample.

Given a pathway-defined gene set *𝒫* ⊆ 𝒱, we aim to develop a statistical framework to test whether genes in *𝒫* exhibit significant enrichment at the extremes of the ranked interactions. The proposed methodology, implemented in GREA, operates through three principal computational phases: (1) Interaction Ranking. We compute entropy-based rank scores for all gene interactions using gene expression profiles, which may depend on phenotype labels when available. (2) Enrichment Quantification. We evaluate the distribution of gene set *𝒫* within the ranked gene interactions using an adaptation of the Kolmogorov-Smirnov (KS) or Recovery-Curve (RC) running sum statistic. (3) Statistical Validation. We determine the statistical significance of the observed enrichment signals by fitting the Gamma distribution to approximate the null distribution, which is generated through pathway-preserving network randomization. In cases where the Gamma distribution provides a poor fit, we instead employ permutation testing to construct the null distribution and compute empirical *p*-values.

#### Calculating Entropy-based Rank Score for Gene Interaction

First, we use NIEE (8) to compute an entropy-based rank score 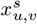 for gene interaction (*u, v*) in sample *s*. This step follows our original NIEE procedure, summarized in the Supplementary Methods.

Given the availability of phenotype labels for the sample-specific data, we derived the phenotype-specific score 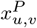 by aligning phenotype labels to samples. We then performed differential gene interaction expression analysis to compare entropy-based scores of gene interaction (*u, v*) across different phenotypic groups. This analysis was conducted using the scanpy package (9) with the Wilcoxon statistical test. Specifically, we compute the log-fold change of 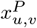 between phenotypic groups to quantify the magnitude and direction of score alterations associated with different conditions. To assess statistical significance, we applied the Wilcoxon rank-sum test, obtaining test statistics that indicate the extent of these differences. The resulting test statistics are ranked in descending order to prioritize gene interactions that exhibit the most significant phenotype-associated entropy shifts. These ranked test statistics serve as input for the subsequent phenotype-based analysis in our algorithmic framework.

#### Calculating Gene Set Enrichment Signal for Ranked Gene Interactions

Recall there are *N* gene interactions, which are ranked based on the sample-specific or phenotype-specific rank scores described in the previous section. For simplicity, let interaction *ℐ*_*i*_ be the *i*-th ranked interaction, and its rank score is *x*_*i*_. Given a pathway-defined gene set *𝒫*, if interaction *ℐ*_*i*_ overlap with *𝒫* (i.e., *ℐ*_*i*_ ∩ *𝒫*≠∅), we define the overlap ratio as 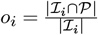, where | · | denotes the cardinality. Clearly, the overlap is determined by the presence of genes in interaction *I*_*i*_ within the gene set: *o*_*i*_ = 0.5 if only one gene is included, and *o*_*i*_ = 1 if both genes are present. Next, we define the hit score *h*_*i*_ as the absolute rank score |*x*_*i*_| of list *ℐ*_*i*_ weighted by the overlap ratio *o*_*i*_, normalized by the sum of the absolute rank scores of all hit lists:

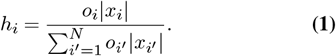

In contrast, if neither gene from interaction *ℐ*_*i*_ is in the gene set, *𝒫* (i.e., *ℐ*_*i*_∩ *𝒫* = ∅), we set the overlap ratio *o*_*i*_ = 0 and define its miss score *m*_*i*_ as the proportion of interactions completely absent from the gene set:

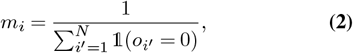

where 𝟙 (·) is an indicator function that equals 1 if the condition is true and 0 otherwise.

For each interaction *ℐ*_*i*_, we perform enrichment analysis using two running sum (RS) statistics: Kolmogorov–Smirnov (KS) (3) approach and recovery curve (RC) approach (10). The RS in the KS-like approach accumulates hit and miss scores from the top-ranked interactions up to a given interaction *ℐ*_*i*_:

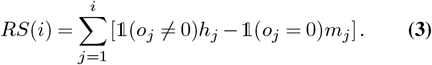

There are two ways to compute the enrichment signal of a given gene set *𝒫* over a list of pre-ranked interactions from the RS. In the first way, the enrichment score (ES) of the entire gene set *𝒫* is then defined as the maximal absolute RS across all interactions:

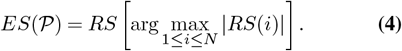

The second way computes the difference between the positive and negative maximum derivatives of scores, which we call the enrichment score difference (ESD):

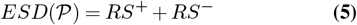

For the full formal definition, see Supplementary Methods.

In the RC approach, the recovery curve is defined as the RS that accumulates only the hit scores from the top-ranked interactions up to a given interaction *ℐ*_*i*_:

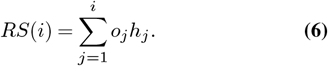

Next, we compute the area under the curve (AUC) as the enrichment signal of a given gene set *𝒫* over a list of pre-ranked interactions from the RS. This AUC value has a range between 0 and 1, and we numerically estimate it using the trapezoidal rule (11):

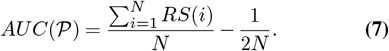

#### Statistical Significance Test

To test whether the pathway gene set *𝒫* is enriched at the extremes of the ranked interactions, we posit the null hypothesis *H*_0_ that *𝒫* is randomly distributed across the ranking (i.e., no top- or bottom-enrichment). We approximate the null and compute *p*-values using two approaches: analytical-based and permutation-based.

In the analytical approach, we randomly shuffle the interaction ranks *M* times (e.g., *M* = 1000) and recalculate the enrichment signal 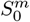 for each shuffle *m*. Next, we approximate the null distribution 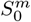 using a bimodal model that separately accounts for positive and negative signals. Specifically, it is modeled as a mixture of two Gamma distributions: one representing all positive signals, *S*^+^, and the other representing all negative signals, *S*^−^:

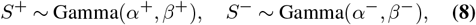

where *α*^+^, *β*^+^ and *α*^−^, *β*^−^ are the shape and scale parameters for the positive and negative Gamma distributions, respectively.

Thus, as detailed in Supplementary Methods, the two-sided *p*-value of an observed enrichment signal *S* is calculated as twice the probability that *S* falls within the bimodal distribution formed by the positive and negative Gamma models:

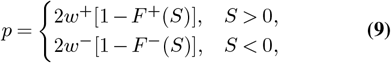

where *F* ^+^(*S*) and *F* ^−^(*S*) denote the cumulative distribution function (CDF) of the positive and negative Gamma distribution, respectively. *w*^+^ and *w*^−^ represent the weight of positive and negative Gamma distributions, respectively, which are determined based on the ratio of positive to negative scores in the null distribution. Specifically, we define 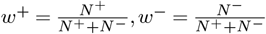, where *N* ^+^ and *N*^−^ denote the number of positive and negative enrichment scores in the null distribution, respectively. This ensures that the weights reflect the ratio of positive to negative scores in the null distribution: *w*^+^ + *w*^−^ = 1.

In contrast to the Kolmogorov–Smirnov (KS) approach, which considers both positive and negative signals, the recovery curve (RC) method focuses solely on positive signals. Following the same procedure, we randomly shuffle the interaction ranks *M* times (e.g., *M* = 1000) and recalculate the enrichment signal 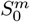 for each shuffle *m*. The null distribution of the enrichment signals is then approximated using a Gamma distribution to account for a positive signal, *S* ∼ Gamma(*α, β*), where *α* and *β* are the shape and scale parameters for the Gamma distribution. Given this model, the statistical significance of an observed enrichment signal *S* is assessed by calculating its position within the estimated distribution. Specifically, the one-side *p*-value is defined as: *p* = 1 − *F* (*S*), where F(S) denotes the CDF of the Gamma Distribution.

However, we found that accurately modeling the RC null with a Gamma distribution is often difficult. Accordingly, we issue warnings for potential misfits and recommend using the permutation-based approach instead (see below).

In the permutation test, the nominal *p*-value is computed by comparing the observed enrichment score *S* to the null distribution of enrichment scores 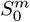 obtained from the permutations. The *p*-value formula is:

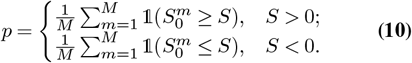

We compare the calculated *p*-value to a pre-defined significance level *α* (commonly set at 0.05) to determine if the gene set enrichment is statistically significant. If *p* ≤ *α*, we reject the null hypothesis, concluding that the gene set *𝒫* is significantly enriched at the top or bottom of the ranked interactions.

Given that multiple gene sets are often tested simultaneously, we apply the False Discovery Rate (FDR) adjustment to control for multiple comparisons with Benjamini-Hochberg (12) and Sidak correction (13) approaches.

Finally, larger gene sets tend to exhibit higher enrichment signals, *S*, purely by chance. To account for variability in gene set size and ensure comparability across different gene sets, we compute the normalized enrichment signal for each gene set, *𝒫*, by dividing the observed enrichment signal by the mean enrichment signal derived from random permutations (null distribution): 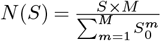.

## Results

### Overview of GREA

We present Gene Interaction Enrichment Analysis (GREA), an enrichment framework that integrates gene–gene regulatory interactions with rank-based scoring to quantify pathway-level differences Figure 1. Designed to address existing limitations with interaction-aware signals, GREA takes three inputs: (1) gene expression, (2) phenotype labels (e.g., case/control), and (3) a gene set library (e.g., KEGG, GO) (Figure 1A).

**Fig. 1.**
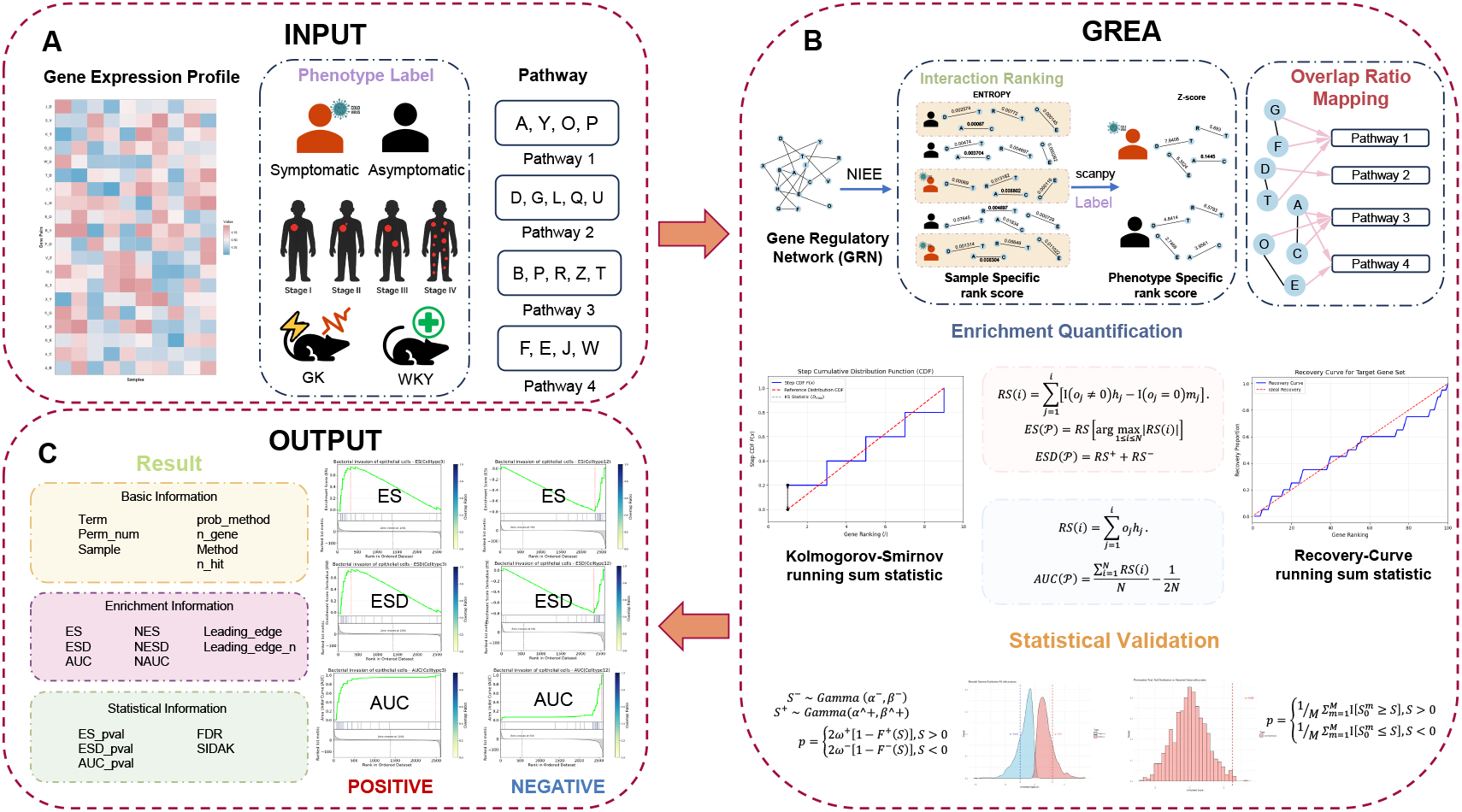
Graphic overview of GREA algorithm. (A) The input for the GREA algorithm is gene expression profiles, phenotype labels (such as symptomatic and asymptomatic), and pathway information. (B) The process of analyzing gene interactions with GREA. The process begins with Interaction Ranking based on gene interaction expression values, yielding sample-specific scores. These scores are then combined with phenotype labels to generate phenotype-specific scores. Overlap ratio mapping illustrates the relationships between different pathways. Then, enrichment quantification employs statistical methods such as the Kolmogorov-Smirnov running sum statistic and the Recovery-Curve running sum statistic to evaluate gene set enrichment. Finally, statistical validation is performed using Gamma distributions and permutation testing to derive *p*-values and determine the significance of the results. (C) The outputs for the algorithm are tabular files containing metrics quantifying the degree of enrichment for specific gene sets. The output also includes visualization plots of ES, ESD, and AUC.

Figure 1B illustrates the core algorithm of GREA. First, we derive sample-specific interaction scores 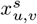 from a gene regulatory network (GRN) using an entropy-based method. These sample-specific interactions are transformed into phenotype-specific ranking score 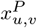 for each gene interaction with the differential test. GREA computes three enrichment signals: (1) Enrichment Score (ES) derived from the Kolmogorov–Smirnov statistic; (2) Enrichment Score Difference (ESD), which quantifies the maximum difference between positive ES and negative ES; (3) Area Under the Curve (AUC) of recovery curves that only focus on the hit gene in the predefined gene set library. Here, we propose a continuous overlap ratio metric that quantifies the degree to which a pathway is represented in high-ranking regulatory interactions. This contrasts with traditional binary hit-counting indicators and improves sensitivity for partially active pathways. After getting the enrichment signals, GREA employs both the permutation test and the gamma distribution fitting to evaluate the statistical significance, enabling flexible hypothesis testing across different data sizes and sparsity levels.

GREA outputs a ranked list of enriched pathways for each phenotype, along with *p*-values and essential information. We also provide the visualization for both KS and RC approaches (Figure 1C).

Overall, GREA links interaction-aware ranking to pathway interpretation, enhancing sensitivity and robustness. Subsequent sections benchmark its performance against state-of-the-art methods on real disease datasets.

### Benchemarking: *p*-value Reproducibility Across Enrichment Methods

In Figure 2, we comprehensively evaluate the performance of different enrichment analysis methods across multiple dimensions. Notably, these analyses are based on a combined dataset aggregating results from multiple disease-specific datasets. To provide further specificity, we have also generated individual figures for each dataset separately Supplementary Figures 2-4. This allows for a more detailed examination of method performance within specific biological contexts.

**Fig. 2.**
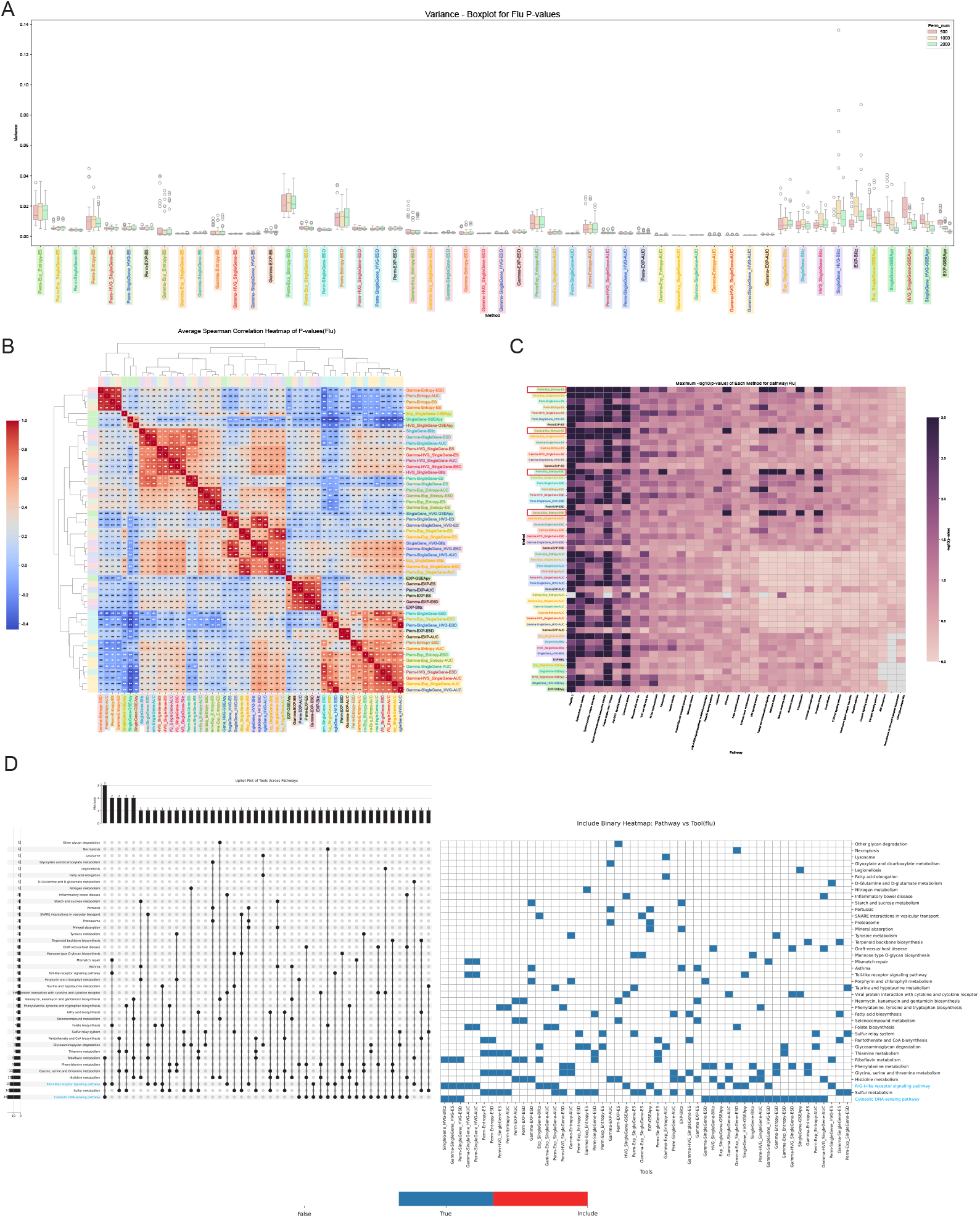
Flu datasets Benchmarking. (A) Box plot comparing the reproducibility of *p*-values across three repeated experiments under different permutations, measured by variance. (B) Heatmap showing the average Spearman correlation of *p*-values between each pair of different methods. Both axes represent different preprocessing methods, and each cell indicates the correlation between the *p*-values obtained from the corresponding pair. (C) Heatmap displaying the maximum − log_10_(*p*-value) obtained by each preprocessing method for its most significantly enriched pathway. The y-axis lists preprocessing methods, and the x-axis shows the corresponding pathways identified by each method. Color intensity reflects the statistical significance, with higher values indicating stronger enrichment. Notably, grey cells represent cases where no significant pathway was detected by the corresponding method. (D) UpSet plot illustrating the overlap of the top 4 significantly enriched pathways identified by different tools on the flu dataset. Each column represents a unique combination of methods, and each row corresponds to a specific enriched pathway. The bar chart indicates the number of pathways jointly identified by each method combination, while the heatmap shows which pathways were identified by which method combinations.

In Figure 2A, we present a comparative boxplot of the variance in enrichment *p*-value estimation reproducibility across all disease datasets for different preprocessing methods and enrichment strategies. The y-axis represents the variance of *p*-values in repeating three times, while the x-axis lists the combinations of preprocessing pipelines, enrichment methods, and statistical models. Each boxplot summarizes the distribution of variances under three different numbers of permutations (500, 1,000, and 2,000), shown in red, beige, and green, respectively.

On the left and middle parts of the figure, we observe results from our method GREA, under different statistical frameworks: permutation testing (Perm), gamma distribution fitting (Gamma), and hybrid approaches like KS and AUC-based scoring. In general, GREA-based methods show consistently lower variance across, with the minimum value in variance being 0.000405 compared to the alternative methods shown on the far right: blitzGSEA and GSEApy, with the minimum variance of 0.001599 and 0.001795, respectively. These two commonly used enrichment tools exhibit higher and more dispersed variance, especially under different pre-processing inputs. Moreover, the blitzGSEA and GSEApy demonstrate more outliers than our GREA methods. This suggests that GREA provides more stable enrichment results, particularly when applying phenotype-specific scores derived from entropy or interaction-based features.

Within the GREA method, the figure highlights that pre-processing methods like Exp_Entropy and Entropy generally lead to higher variance, 0.044748, than other methods. Gamma-based models also appear to reduce variability further in certain cases, like all preprocessing methods in AUC. These internal comparisons clearly show that the gamma distribution fitting adds more stability to *p*-value production than permutation testing.

Same dataset for Figure 2B, we want to evaluate the similarity between our method and other methods in past research. Spearman correlation (14) indicates that the two methods assign similar levels of significance to the same sets of pathways across diverse biological contexts.

Similar to initial expectations, it is worth noting that our unique part of analyzing gene interactions in GREA is not very similar to the past two methods, both of which are *ρ* < 0.5. The results in blitz are highly connected with our enrichment in ES, ESD in gamma fitting, and AUC in permutation testing. By contrast, methods based on the GSEApy enrichment method (e.g., SingleGene-GSEApy, HVG_SingleGene-GSEApy) exhibit low or even negative correlation with nearly all other methods (*ρ* < 0.3), including both gamma- and permutation-based GREA configurations. This indicates that GSEApy scoring yields a different enrichment rank.

Within our GREA methods, a clear block of high internal similarity is observed among GREA configurations between AUC in gamma fitting and ESD in permutation testing, the same as in AUC in permutation testing and ESD in gamma fitting. The detailed results of the variance, mean, and sample number for different repetitions, permutations, and methods are provided in the Supplementary Table 1.

As shown in Figure 2C, the performance of different enrichment analysis methods varies significantly in terms of pathway detection sensitivity, as measured by the maximum − log_10_(*p*-value) across pathways. Notably, past methods blitzGSEA and GSEApy, particularly those using HVG- or SingleGene-based gene sets, exhibit poor performance on the right-hand side of the heatmap, where a substantial number of pathways show extremely low significance levels or are entirely undetected (grey or pale pink regions). This indicates a limited capacity of these methods to identify relevant biological pathways under these settings.

In contrast, our proposed method, GREA, demonstrates a clear advantage in both sensitivity and coverage. When combined with the EXP_Entropy gene selection strategy and either the ES or ESD enrichment approach (Perm-Exp_Entropy-ES, Gamma-Exp_Entropy-ES, Perm-Exp_Entropy-ESD, Gamma-Exp_Entropy-ESD), GREA consistently identifies more pathways with significantly higher −log_10_(*p*-value), especially in the pathway Ribosome, Retrograde endocannabinoid signaling, Parkinson disease, Thermogenesis and Huntington disease. These results highlight the effectiveness of incorporating interaction gene pair information in enhancing signal detection and biological relevance. The ability of GREA to capture additional significant pathways not detected by other methods suggests that its design better reflects the underlying regulatory or functional relationships in the data. Overall, this comparison underscores the robustness and improved interpretability of GREA in complex pathway enrichment scenarios. Detailed information on the minimum *p*-values, maximum −log_10_(*p*-value), and corresponding pathways for each method can be found in Supplementary Table 2.

Additionally, to validate the theoretical behaviour of KS and RC, we conducted extreme scenario analyses where all genes in the ranked gene list are either completely included (all-hit) or completely absent (non-hit) in the pathway library, as illustrated in Supplementary Figure 1. These boundary condition tests demonstrate the expected mathematical properties.

### Case Study: Respiratory Viral Infection Analysis

To evaluate the biological applicability of our enrichment analysis method, we conducted a case study using publicly available transcriptomic datasets derived from seven controlled human viral challenge studies. We included two public human influenza challenge datasets, GSE52428 (15) and GSE73072 (16). For the former one, in total, 41 volunteers participated in these two studies, of whom 18 developed symptomatic infection and were assigned to the infection group, while 23 remained asymptomatic or had no detectable infection and were treated as controls. For later ones, these studies involved a total of 148 healthy adult volunteers who were experimentally inoculated with one of four respiratory viruses: Influenza A/H1N1, Influenza A/H3N2, Respiratory Syncytial Virus (RSV), or Human Rhinovirus (HRV). We integrated the dataset by viral subtypes and applied our enrichment analysis approach to identify the common pathways for influenza. We extracted the top four significantly enrichment signals from each method—GREA, as well as two established tools: blitzGSEA and GSEApy—to facilitate a comparative evaluation. It is worth noting that by applying at least three different preprocessing strategies and combinations of the GREA method, the Influenza A pathway (17, 18)was consistently identified across multiple datasets, suggesting the realizability and biological relevance of our method in response to viral infection. Moreover, when the enrichment signal threshold of the pathway was raised to the top 10, almost all combinations were able to identify Influenza A as a significantly enriched pathway, which was highly consistent with the results obtained by existing methods.

Beyond the expected Influenza A pathway, as shown in Figure 2D, our method also revealed strong and useful enrichment signals in two additional pathways with known relevance to viral infection: the Cytosolic DNA-sensing pathway (hsa04623) (19) and the RIG-I-like receptor signalling pathway (hsa04622) (17, 20). These pathways are critical for host defence against RNA viruses, such as Influenza A, as they mediate viral nucleic acid recognition and activate downstream type I interferon responses. These observations suggest that our method not only detects classical pathway enrichment but also captures strong connection interaction-level signals.

### Case Study: Cancer Pathway Enrichment Analysis

We systematically evaluated five cancer types—colon adenocarcinoma (COAD), kidney renal clear cell carcinoma (KIRC), lung adenocarcinoma (LUAD), stomach adenocarcinoma (STAD), and thyroid carcinoma (THCA) (21)—to benchmark the enrichment performance of our method against two existing tools: blitzGSEA and GSEApy, through the *p*-value, and find the unique pathway to illustrate the new biological meanings by the top 4 enrichment signals. For each dataset, we systematically investigate how different enrichment analysis methods perform when coupled with diverse preprocessing approaches (Figure 3 and Supplementary Figures 5–7). Figure 3A shows p-value variance. The left/center panels report GREA hybrid metrics across preprocessing methods, statistical paradigms (Perm, Gamma), and scoring functions (KS, AUC). GREA variants consistently yield lower variance than benchmarks on the right (blitzGSEA min = 0.006412; GSEApy min = 0.005662 vs GREA min = 0.001421), with benchmarks exhibiting wider spreads and more outliers across preprocessing settings. Overall, GREA produces more stable enrichment results, likely due to its gene-interaction modeling. Within GREA, some preprocessing choices markedly increase variance. In particular, Exp_Entropy and Entropy yield higher variability with permutation-based enrichment, reaching up to 0.003915. By contrast, gamma-based modeling further suppresses variance—especially with AUC-based scoring—highlighting that gamma fitting stabilizes p-value estimation compared with permutation alone.

**Fig. 3.**
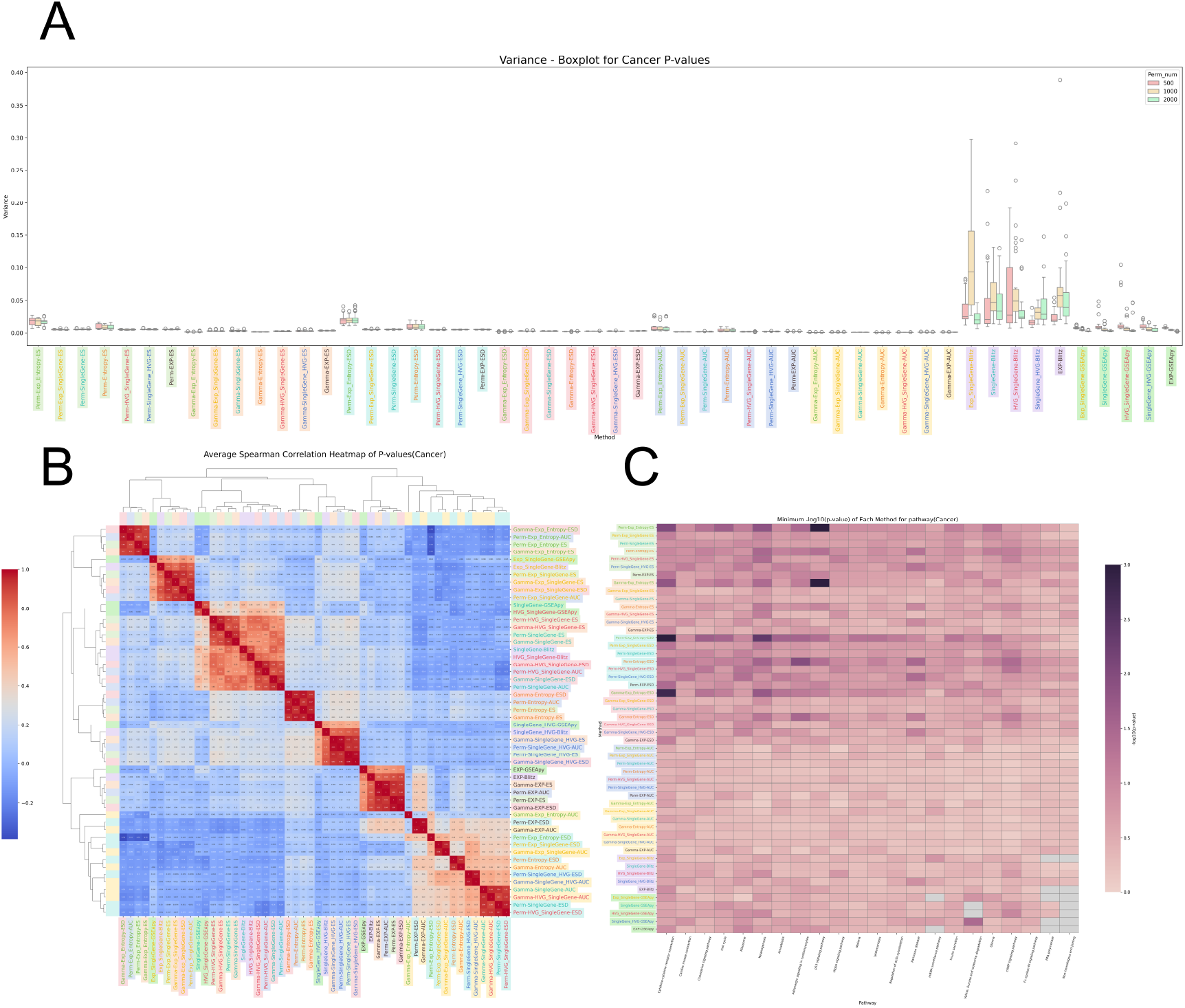
Cancer datasets benchmarking. (A) Comparison of *p*-value reproducibility between different permutations in three-fold replication through calculating the variance in the boxplot. The y-axis denotes the dispersion of *p*-values obtained from three repeated runs, whereas the x-axis displays distinct combinations of preprocessing pipelines, enrichment procedures, and statistical evaluation models. Variance distributions for three separate permutation depths (500, 1,000, and 2,000) are differentiated by red, beige, and green boxplots, respectively. (B) The average Spearman correlation coefficients of pathway enrichment *p*-values across pairs of preprocessing methods applied to different cancer datasets are shown in this heatmap, where both axes represent the different preprocessing strategies and each cell indicates the degree of correlation between them. (C) The maximum − log_10_(*p*-value) for the most significantly enriched pathway identified by each preprocessing method in different cancer datasets is illustrated in this heatmap, with the y-axis listing the preprocessing approaches and the x-axis showing their top-ranked pathways. The gradient of color denoted the statistical significance, with increased darkness indicating greater significance and stronger pathway enrichment. In contrast, cells shaded in gray denote that the corresponding preprocessing method did not yield any significantly enriched pathway.

In Figure 3B, we assess pairwise concordance across enrichment strategies and preprocessing pipelines for cancer datasets using Spearman correlations of pathway-level *p*-value ranks. The hierarchically clustered heatmap reveals coherent method groupings. GREA variants that leverage interaction-derived features cluster together, showing moderate-to-high intra-group correlations. Strong associations are seen between AUC with Gamma modeling and ESD with permutation testing, indicating internal consistency within GREA. Dark blocks in the upper-left quadrant indicate that these GREA configurations assign similar pathway significance (higher correlations). In contrast, GSEApy methods—especially with SingleGene or HVG gene sets— show much lower or even negative correlations (*ρ* < 0.3) relative to most GREA and blitz settings, implying divergent pathway rankings under cancer-specific preprocessing. blitzGSEA exhibits moderate concordance with selected GREA variants, particularly for ES and ESD under both Gamma-modeled and permutation-based inference. Full correlation matrices and summary statistics (including variance and permutation replicate counts) are provided in Supplementary Table 3.

Figure 3C compares pathway detection sensitivity using the maximum −log_10_(*p*-value) achieved per pathway. per pathway. Although overall performance is similar, blitzGSEA and GSEApy miss certain pathways under specific preprocessing. For example, Non-homologous end-joining is never detected by GSEApy, and RNA polymerase is consistently missed by both GSEApy and blitzGSEA with Exp_SingleGene or Exp preprocessing (gray cells).

By contrast, GREA paired with EXP_Entropy and ES/ESD modes (Perm-Exp_Entropy-ES, Gamma-Exp_Entropy-ES, Perm-Exp_Entropy-ESD, Gamma-Exp_Entropy-ESD) yields stronger signals for pathways such as Cytokine–cytokine receptor interaction and p53 signaling (dark regions, upper left), indicating enhanced sensitivity and interpretability from interaction-aware modeling. Detailed minima *p*-values, peak −log_10_(*p*-value), and pathway lists are provided in Supplementary Table 4.

Moreover, we performed cancer-stage-specific enrichment analyses using interaction-derived entropy scores and compared against expression-based methods, summarizing shared and unique pathways in Supplementary Figures 8-9. For COAD (Supplementary Figure 8A-D), JAK-STAT signaling (22, 23) common to all methods in stage IIa, and Wnt signaling (24) in stage IV. Unique to our method: Base Excision Repair (25, 26) and Fructose/Mannose Metabolism (27, 28) in stage IIa, and Non-Homologous End Joining (NHEJ) (29) in stage IV. For KIRC (Supplementary Figure 8E-H), PPAR signaling (30) was shared in stage III; Cell Cycle and p53 signaling (31) were common in stage IV. Our method uniquely identified Fatty Acid Biosynthesis (32, 33) and Mismatch Repair (34) in stage IV. For LUAD (Supplementary Figure 8I-L), ECM–receptor interaction (35, 36) was shared in stage IIb; DNA replication (37) and Fanconi anemia (38) in stage IV. Unique to our method: Ubiquitin-mediated proteolysis (39) and Ribosome biogenesis in eukaryotes (40) in stage IIb; Pentose Phosphate Pathway, NHEJ, and Nucleotide Excision Repair (41, 42) in stage IV. For STAD (Supplementary Figure 9A-D), Wnt signaling (43) and Proteoglycans in cancer (44) were shared in stage IIIa; Hematopoietic cell lineage (45) in stage IV. Unique to our method: Other types of O-glycan biosynthesis (46, 47) and Allograft Rejection (48, 49) in stage IIIa; Adherens Junction (50) and Nucleotide Excision Repair (51) in stage IV. For THCA (Supplementary Figure 9E-H), ECM–receptor interaction and Cytokine–cytokine receptor interaction (52–54) were shared in stage III; ECM–receptor interaction remained enriched in stage IVa. Unique to our method: Hedgehog signaling (55, 56) and NHEJ (57) in stage III; Cholesterol Metabolism (58) and Mitophagy (59) in stage IVa.

Across cancers, recurrent pathways detected by all methods included JAK–STAT (COAD), PPAR (KIRC), ECM–receptor interaction (LUAD, THCA), and Wnt signaling (COAD, STAD), indicating consistent activation of key regulators. In contrast, the unique pathways above arose exclusively from interaction-entropy scoring and were not detected by blitzGSEA or GSEApy, underscoring our method’s sensitivity to interaction-centric signals. Full results are in Supplementary Table 5.

## Discussion

In this study, we introduced GREA, a novel framework that integrates gene regulatory interactions into pathway library enrichment analysis. Our results demonstrate that GREA offers several key advantages over traditional GSEA methods, particularly in its ability to capture subtle but biologically meaningful signals in gene interactions.

Through *p*-value benchmarking experiments, we showed that GREA produces more stable and reproducible results compared to past enrichment tools like blitzGSEA (4) and GSEApy (7). The incorporation of continuous overlap ratios and different statistical testing approaches (permutation testing and gamma distribution fitting) significantly enhances the robustness of enrichment signal detection. Notably, GREA consistently identified more significant pathways across diverse preprocessing strategies, suggesting its versatility in handling different types of input data.

One of the most important contributions of GREA is its focus on gene interactions rather than individual genes. This approach gave a novel horizon of biological systems, where subtle difference is often found in coordinated interactions between genes. The ability of GREA to detect partial pathway activation and subtle changes in regulatory networks represents a substantial improvement over binary hit-counting methods used in traditional GSEA.

Furthermore, we observed that genes may play different roles in different pathways - for example, Gene Ontology Biological Process (GOBP) pathways focus on the involvement of genes in biological processes, while Gene Ontology Molecular Function (GOMF) pathways emphasize the specific activities performed by genes. In this context, applying gene interaction-based analysis may yield different enrichment results. In future work, we plan to further investigate and compare the performance of GREA across these different pathway types.

Despite these advances, some limitations remain. While the gamma distribution fitting generally improved stability, we observed challenges in accurately modeling the null distribution for recovery curve-based enrichment. Future work should focus on refining these aspects and extending the capabilities of GREA to single-sample analysis, which would be particularly valuable for biological pathway identification applications.

In conclusion, GREA represents a significant step forward in gene set enrichment analysis by explicitly considering gene interactions and employing advanced statistical methodologies. As biological research continues to generate increasingly complex datasets, tools like GREA will be essential for extracting meaningful insights and advancing our understanding of cellular processes.

## Data Availability

The TCGA cancer data are available from https://portal.gdc.cancer.gov/. The GEO influenza data are available under accessions GSE52428 and GSE73072.

## Code Availability

The GREA source code is available at https://github.com/compbioclub/GREA.

## Acknowledgements

We express our gratitude for the support provided by the National Natural Science Foundation of China (No. 32400519 - LXC) and the CityU Tung Biomedical Sciences Centre Project Fund (No. 9609331 - LXC).

## Author Contributions Statement

XYL: Software; Formal analysis; Validation; Visualization; Investigation; Methodology; Writing—original draft; Writing—review and editing. ANJ: Software; Formal analysis; Validation; Visualization; Investigation; Methodology; Writing—original draft; Writing—review and editing. CSL: Formal analysis; Validation; Investigation; Writing—original draft; Writing—review and editing. LXC: Conceptualization; Supervision; Software; Methodology; Writing—original draft; Writing—review and editing.

## Competing Interests

The authors declare no competing interests.

## Notes

https://github.com/compbioclub/GREA

### Competing Interest Statement

The authors have declared no competing interest.

### Summary of Updates

Add real-world case studies using viral infection and cancer datasets.

