## Supplementary File for "Gene interaction enrichment analysis for transcriptomic data with GREA"

8 (A, C, E, G, I, H) UpSet plots illustrate the overlap of the top 4 significantly enriched pathway combinations identified in COAD stage IIA, COAD stage IV, KIRC stage III, KIRC stage IV, LUAD stage IIA, and LUAD stage IV. The accompanying heatmaps display the specific methods associated with each pathway combination. Red highlights indicate combinations that include all three methods: GREA, blitzGSEA, and GSEApy. (B, D, F, H, J, L) UpSet plots show the top 4 significantly enriched pathway combination overlaps for COAD stage IIA, COAD stage IV, KIRC stage III, KIRC stage IV, LUAD stage IIA, and LUAD stage IV. In the corresponding heatmaps, red highlights mark pathway combinations that are identified solely by GREA, without contributions from blitzGSEA or GSEApy. . . . 14

9 (A, C, E, G) UpSet plots illustrate the overlap of the top 4 significantly enriched pathway combinations identified in STAD stage IIIA, STAD stage IV, THCA stage III, and THCA stage IVA. The accompanying heatmaps display the specific methods associated with each pathway combination. Red highlights indicate combinations that include all three methods: GREA, blitzGSEA, and GSEApy. (B, D, F, H) UpSet plots show the top 4 significantly enriched pathway combination overlaps for STAD stage IIIA, STAD stage IV, THCA stage III, and THCA stage IVA. In the corresponding heatmaps, red highlights mark pathway combinations that are identified solely by GREA, without contributions from blitzGSEA or GSEApy. . . . . . . . . . . . . . . . . . . . . . 15

### 1 Supplementary Methods

#### 1.1 Calculating Entropy-based Rank Score for Gene Interaction

We leverage the NIEE algorithm [1] to calculate an entropy-based rank score $x^s_{u,v}$ for gene interaction $(u, v)$ in sample $s$. First, the top 5,000 highly variable genes (HVGs) are screened from the input dataset by the scanpy toolkit with parameter `flavor=seurat_v3` [2, 3]. StringDB Protein-Protein Interaction Networks (`score>850`) are then employed as the background GRN to form the graph $G = (\mathcal{V}, \mathcal{E})$ [4].

##### 1.1.1 Constructing reference network

The reference network is constructed using all reference samples $\mathcal{R}$, usually representing relatively healthy or control samples. For each gene $u$, its reference entropy, $H_u(\mathcal{R})$, is calculated as:

$$H_u(\mathcal{R}) = -\frac{1}{\log L} \sum_{w=1}^{L} p_{u,w}(\mathcal{R}) \log p_{u,w}(\mathcal{R}), \tag{1}$$

where $w$ represents the first-order neighbor genes of gene $u$, and $L$ is the number of neighbors for gene $u$. $p_{u,w}(\mathcal{R})$ is the probability of the interaction between gene $u$ and $w$, defined as:

$$p_{u,w}(\mathcal{R}) = \frac{|PCC_{u,w}(\mathcal{R})|}{\sum_{w=1}^{L} |PCC_{u,w}(\mathcal{R})|}, \tag{2}$$

where $PCC_{u,w}(\mathcal{R})$ is the Pearson correlation coefficient between gene $u$ and $w$ in $\mathcal{R}$ reference samples.

The reference entropy $H_{u,v}(\mathcal{R})$ for gene interaction $(u, v)$ can be calculated as:

$$H_{u,v}(\mathcal{R}) = \frac{H_u(\mathcal{R}) + H_v(\mathcal{R})}{2}. \tag{3}$$

Similarly, the reference standard deviation $SD_{u,v}(\mathcal{R})$ for gene interaction $(u, v)$ is:

$$SD_{u,v}(\mathcal{R}) = \frac{SD_u(\mathcal{R}) + SD_v(\mathcal{R})}{2}, \tag{4}$$

where $SD_u(\mathcal{R})$ is the standard deviation for gene $u$ across the reference samples $\mathcal{R}$.

##### 1.1.2 Constructing perturbation network

After constructing the reference network, a sample $s$ is added to the reference samples $\mathcal{R}$ to form the sample-specific network $\mathcal{R}^s$. The sample-specific entropy $H_{u,v}(\mathcal{R}^s)$ and the standard deviation $SD_{u,v}(\mathcal{R}^s)$ for gene interaction $(u, v)$ are then calculated following the same procedure as in the reference network.

##### 1.1.3 Calculating the entropy of edge

To quantify the fluctuations caused by sample $s$, we can calculate the sample-specific entropy-based rank score $x^s_{u,v}$ for gene interaction $(u, v)$ as:

$$\begin{aligned} x^s_{u,v} &= |H_{u,v}(\mathcal{R}^s) - H_{u,v}(\mathcal{R})| \\ &\cdot |SD_{u,v}(\mathcal{R}^s) - SD_{u,v}(\mathcal{R})| \end{aligned} \tag{5}$$

#### 1.2 Detailed Formula for ESD

This section presents a detailed definition of the enrichment score difference (ESD). The ESD is a measure that combines both the maximum positive and negative deviations of the scores. The ESD formula is defined as follows:

$$ESD(\mathcal{P}) = \begin{cases} RS\left[\arg\max_{1\le i\le N} RS(i)\right] + RS\left[\arg\min_{1\le i\le N} RS(i)\right], & \text{if } \exists i \text{ such that } RS(i) > 0 \text{ and } RS(i) < 0 \\ RS\left[\arg\max_{1\le i\le N} RS(i)\right], & \text{if } RS(i) \ge 0 \text{ for all } i \\ RS\left[\arg\min_{1\le i\le N} RS(i)\right], & \text{if } RS(i) \le 0 \text{ for all } i \end{cases} \tag{6}$$

#### 1.3 Modeling enrichment signal using bimodal distribution

In the analytical approach, we approximate the null distribution $S_0^m$ using a bimodal model that separately accounts for positive and negative signals. Specifically, it is modelled as a mixture of two Gamma distributions: one representing all positive signals, $S^+$, and the other representing all negative signals, $S^-$:

$$\begin{aligned} S^+ &\sim \text{Gamma}(\alpha^+, \beta^+), \\ S^- &\sim \text{Gamma}(\alpha^-, \beta^-), \end{aligned} \tag{7}$$

where $\alpha^+$, $\beta^+$ and $\alpha^-$, $\beta^-$ are the shape and scale parameters for the positive and negative Gamma distributions, respectively.

##### 1.3.1 Mixture model formulation

Let $w^+$ and $w^-$ denote the weights of the positive and negative components, with

$$w^+ + w^- = 1, \quad w^+, w^- \geq 0.$$

The probability density function (PDF) of the null distribution $f_S(s)$ is then:

$$f_S(s) = \begin{cases} w^+ f_{S^+}(s), & s > 0, \\ w^- f_{S^-}(-s), & s < 0, \end{cases} \tag{8}$$

where $f_{S^+}$ and $f_{S^-}$ are the Gamma PDFs:

$$f_{S^\pm}(x) = \frac{x^{\alpha^\pm - 1} e^{-x/\beta^\pm}}{\beta^{\pm\alpha^\pm} \Gamma(\alpha^\pm)}, \quad x > 0. \tag{9}$$

##### 1.3.2 Cumulative distribution functions (CDFs)

The corresponding CDFs for $S^+$ and $S^-$ are

$$\begin{aligned} F^+(S) = P(S^+ \leq s) &= \frac{1}{\Gamma(\alpha^+)} \gamma \left( \alpha^+, \frac{s}{\beta^+} \right), \quad s > 0, \\ F^-(S) = P(S^- \leq s) &= \frac{1}{\Gamma(\alpha^-)} \gamma \left( \alpha^-, \frac{s}{\beta^-} \right), \quad s > 0, \end{aligned} \tag{10}$$

where $\Gamma(\cdot)$ denotes the Gamma function, and $\gamma(\cdot, \cdot)$ is the lower incomplete Gamma function [5].

##### 1.3.3 Deriving the $p$-value

Given an observed enrichment score $S = s$, the one-sided $p$-value is defined as the probability under the null to observe a signal as or more extreme than $s$ in the corresponding tail:

- If $s > 0$, the $p$-value is the weighted upper tail probability of the positive Gamma distribution:

$$p' = w^+ P(S^+ \geq s) = w^+ \left[ 1 - F^+(s) \right]. \tag{11}$$

- If $s < 0$, since the negative signals are modeled via the distribution of $-S$, the $p$-value is:

$$p' = w^- P(S^- \geq -s) = w^- \left[ 1 - F^-(-s) \right]. \tag{12}$$

##### 1.3.4 Two-sided $p$-value

A two-sided $p$-value accounts for both extreme positive and extreme negative signals. To capture significance from both directions, the final two-sided $p$-value is:

$$p = 2p'. \tag{13}$$

This ensures the probability of an observed enrichment signal at least as extreme as $s$ in the positive or negative direction, and its statistical significance is assessed symmetrically or approximately balanced between tails.

##### 1.3.5 Parameter estimation

Gamma parameters were estimated using maximum likelihood estimation (MLE) with location fixed at zero, implemented via the `scipy.stats.gamma.fit` routine.

#### 1.4 Multiple Testing Adjustment

The Benjamini-Hochberg method [6] controls the FDR by ranking original $p$-values and applying a step-by-step procedure. Given $m$ as the total number of hypothesis tests, with $p_{(1)} \leq p_{(2)} \leq ... \leq p_{(m)}$ representing $p$-values arranged in ascending order, the corrected $p$-value for the $i$-th test was calculated as:

$$\hat{p}_{(i)} = \min(p_{(i)} \times \frac{m}{i}, 1) \tag{14}$$

This method ensures that the expected proportion of false positives among rejected hypotheses is controlled at level $\alpha$.

We also applied the more conservative Sidak correction [7], which controls the family-wise error rate (FWER). For $m$ independent tests, the corrected $p$-value was calculated as:

$$\hat{p}_i = 1 - (1 - p_i)^m \tag{15}$$

where $p_i$ is the original $p$-value for the $i$-th test. This correction guarantees that the probability of making one or more Type I errors across all tests is less than $\alpha$.

#### 1.5 Experiment setting

##### 1.5.1 Dataset Description

To evaluate the performance of GREA, we conducted a series of benchmarking experiments using publicly available transcriptomic datasets sourced from the NCBI Gene Expression Omnibus (GEO) database (`https://www.ncbi.nlm.nih.gov/geo/`). Specifically, we selected two datasets, GSE52428 [8] and GSE73072 [9], both of which profile host gene expression responses to experimental respiratory viral infections. GSE52428 contains whole blood transcriptional profiles from human subjects who were challenged with either influenza H1N1 or H3N2 viruses. GSE73072 comprises a broader range of respiratory virus challenges, including influenza A subtypes (H1N1 and H3N2), human rhinovirus (HRV), and respiratory syncytial virus (RSV).

In addition to viral infection datasets, we also applied GREA to several cancer transcriptomic datasets obtained from The Cancer Genome Atlas (TCGA) project (`http://cancergenome.nih.gov/` ). Specifically, we selected datasets covering five major cancer types: colon adenocarcinoma (COAD), kidney renal clear cell carcinoma (KIRC), lung adenocarcinoma (LUAD), stomach adenocarcinoma (STAD), and thyroid carcinoma (THCA). These datasets provide comprehensive gene expression profiles from tumor samples, allowing us to evaluate the performance of GREA in detecting pathway enrichment in the context of diverse cancer types.

Detailed application of the GREA algorithm on these datasets enables a comprehensive assessment of its capability to capture biologically meaningful enrichment signals. In the following, we present the complete analytical workflow and key findings obtained from applying GREA to the selected viral infection datasets and benchmarking between GREA and past enrichment methods.

##### 1.5.2 $p$-value Benchmarking

The primary goal of this $p$-value benchmark analysis is to evaluate and compare the reproducibility and similarity of different enrichment methods. To achieve this, we designed multiple preprocessing strategies that generate phenotype-specific scores by ranking z-scores derived from differential tests. We used gene expression or entropy-based interaction data to construct the input matrix for all methods. The preprocessing methods are summarised as follows: All preprocessing strategies share common characteristics, including the selection of 5,000 highly variable features (either genes or gene interactions), the transformation from sample-specific scores to phenotype-specific scores, and the integration of gene expression data with entropy-based interaction information in various combinations.

The specific preprocessing methods include: *Exp_Entropy* involves selecting the top 5,000 highly variable genes (HVGs) from expression data, identifying these genes within entropy-based gene interactions with sample-specific scores, and then transforming them into phenotype-specific scores. This method is used for internal evaluation of our approach. *Exp_SingleGene*also starts by selecting 5,000 HVGs from expression data and identifying corresponding entropy gene interactions, but then decomposes each gene pair into individual genes by averaging their sample-specific scores before converting to phenotype-specific scores. *SingleGene* first selects 5,000 highly variable entropy gene interactions, transforms their sample-specific scores to phenotype-specific scores, and then averages the phenotype-specific scores for each gene to derive single-gene scores. *Entropy*, another internal baseline, selects 5,000 highly variable entropy gene interactions and directly transforms sample-specific scores into phenotype-specific scores without further decomposition. *HVG_SingleGene* begins by selecting 5,000 highly variable entropy gene interactions, decomposes them into single genes by averaging sample-specific scores, and then transforms these into phenotype-specific scores. *SingleGene_HVG* first decomposes gene interactions into single genes with averaging sample-specific value, then selects 5,000 HVGs based on the resulting scores, followed by phenotype-specific transformation. Finally, *EXP* selects 5,000 HVGs directly from expression data and transforms their sample-specific scores into phenotype-specific scores. These diverse preprocessing schemes enable a comprehensive evaluation of enrichment methods under various input conditions.

In the first experiment for $p$-value, we used the Enrichr [10] gene set library created from KEGG [11] using pre-rank in GSEApy [12], blitzGSEA [13], and GREA with 500, 1,000, and 2,000 permutations applied to different datasets related to respiratory viral infections in different preprocessing methods. We examined the consistency of $p$-values across three duplications with the same number of permutations. After calculating the Spearman correlation (corr) [14] between $p$-values, which means the similarity of results, we set the variances as $1 - corr$ and recorded based on the number of permutations for GREA on ES, GREA on ESD, GREA on AUC, blitzGSEA [13], and GSEApy [12]. If the correlation is high (close to 1), the error approaches 0, indicating stable results. If the correlation is low, the error approaches 1, indicating unstable results.

For the second experiment, similar to the setting in the first experiment, we computed pairwise Spearman rank correlations across all method combinations to assess how consistently they ranked pathway significance. This process was repeated for all (repetition time and Permutation number) combinations, and the resulting correlation matrices were averaged to produce a final mean Spearman correlation matrix.

To find the minimum $p$-value pathway under different methods, we set up the third experiment. The color scale represents the $-\log_{10}(p\text{-value})$ values, ranging from light pink ($-\log_{10}(p\text{-value}) \sim 0$) to dark purple ($-\log_{10}(p\text{-value}) \sim 3$). Grey blocks indicate that a particular method did not identify a significant pathway.

These diverse preprocessing approaches enable a comprehensive evaluation of enrichment methods under various input conditions.

### 2 Supplementary Figures

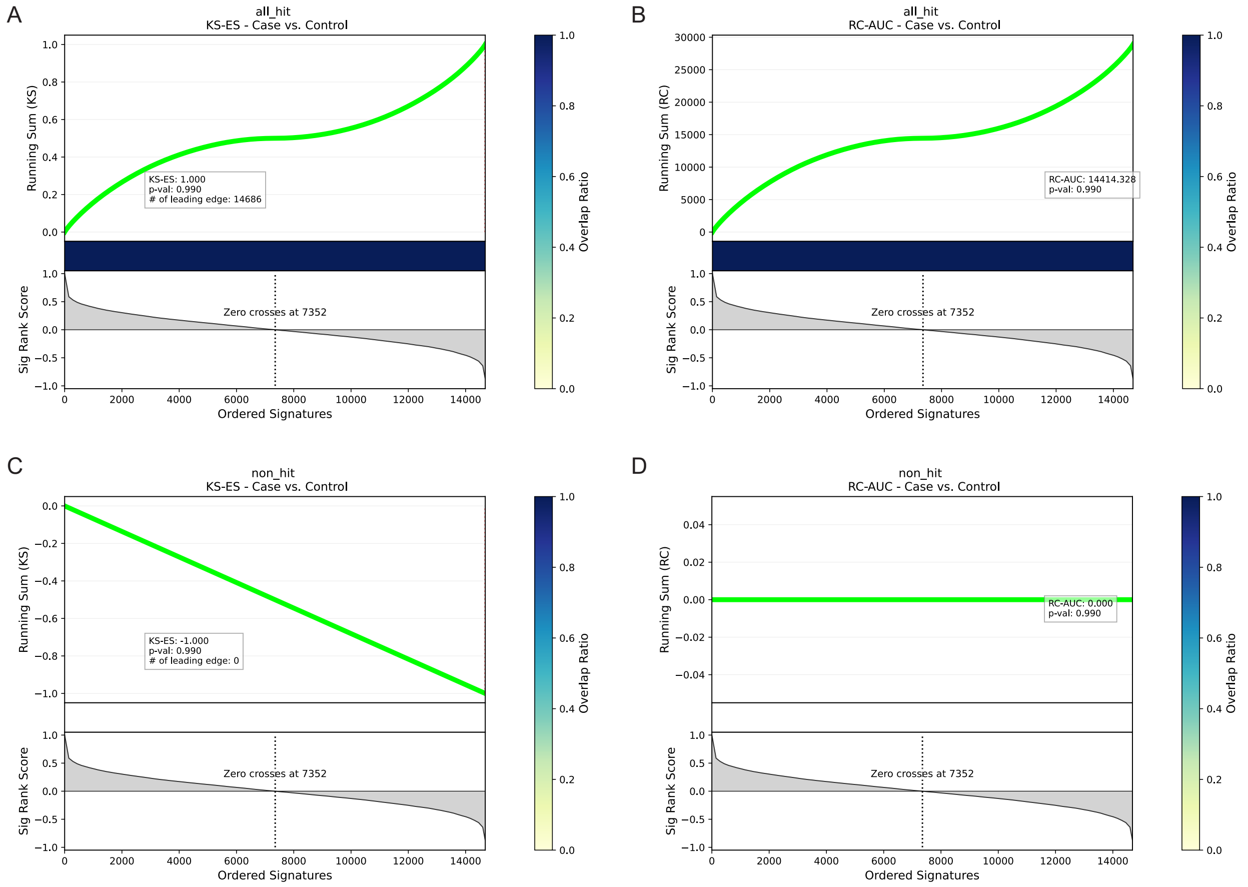

Supplementary Figure 1: Gene set enrichment analysis using Kolmogorov-Smirnov (KS) and Recovery Curve-Area Under Curve (RC-AUC) methods under extreme scenarios.
(A)When all genes in the ranked list are contained in the pathway, the running sum monotonically increases, yielding the maximum enrichment score (ES = 1.0) with all 14,686 genes contributing to the leading edge ($p$-value = 0.990). (B)RC-AUC enrichment analysis under the same all hit scenarios, showing positive enrichment with an AUC value of 14414.328 ($p$-value 0.990). (C)When no genes in the gene set are present in the pathway, the running enrichment score monotonically decreases to the minimum value (ES = -1.0), with no leading-edge genes ($p$-value=0.990). (D)RC-AUC enrichment analysis under the non-hit scenarios, showing no enrichment with an AUC value of 0.000 ($p$-value=0.990).

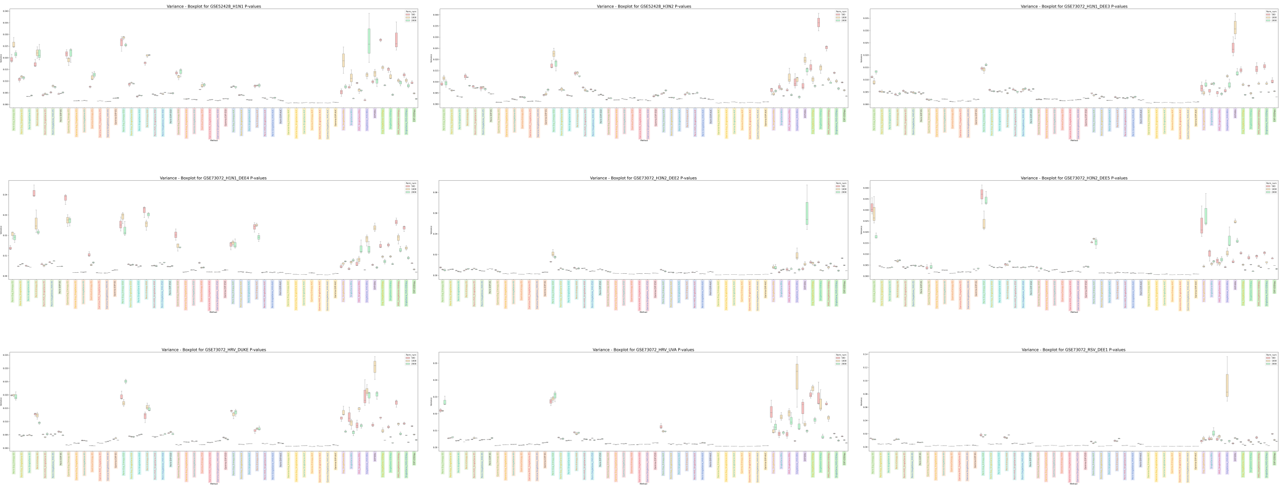

Supplementary Figure 2: $p$-value variance Comparison between different methods in three repeated experiments with 500, 1000, and 2000 permutations for different disease datasets.

The supplementary figure provides a detailed visualization of the reproducibility of $p$-values across individual datasets analyzed under different permutations. Each boxplot corresponds to a flu dataset within the combined dataset in the main text, including datasets GSE52428 with either influenza H1N1 or H3N2 viruses and GSE73072 with influenza A subtypes (H1N1 and H3N2), human rhinovirus (HRV), and respiratory syncytial virus (RSV).

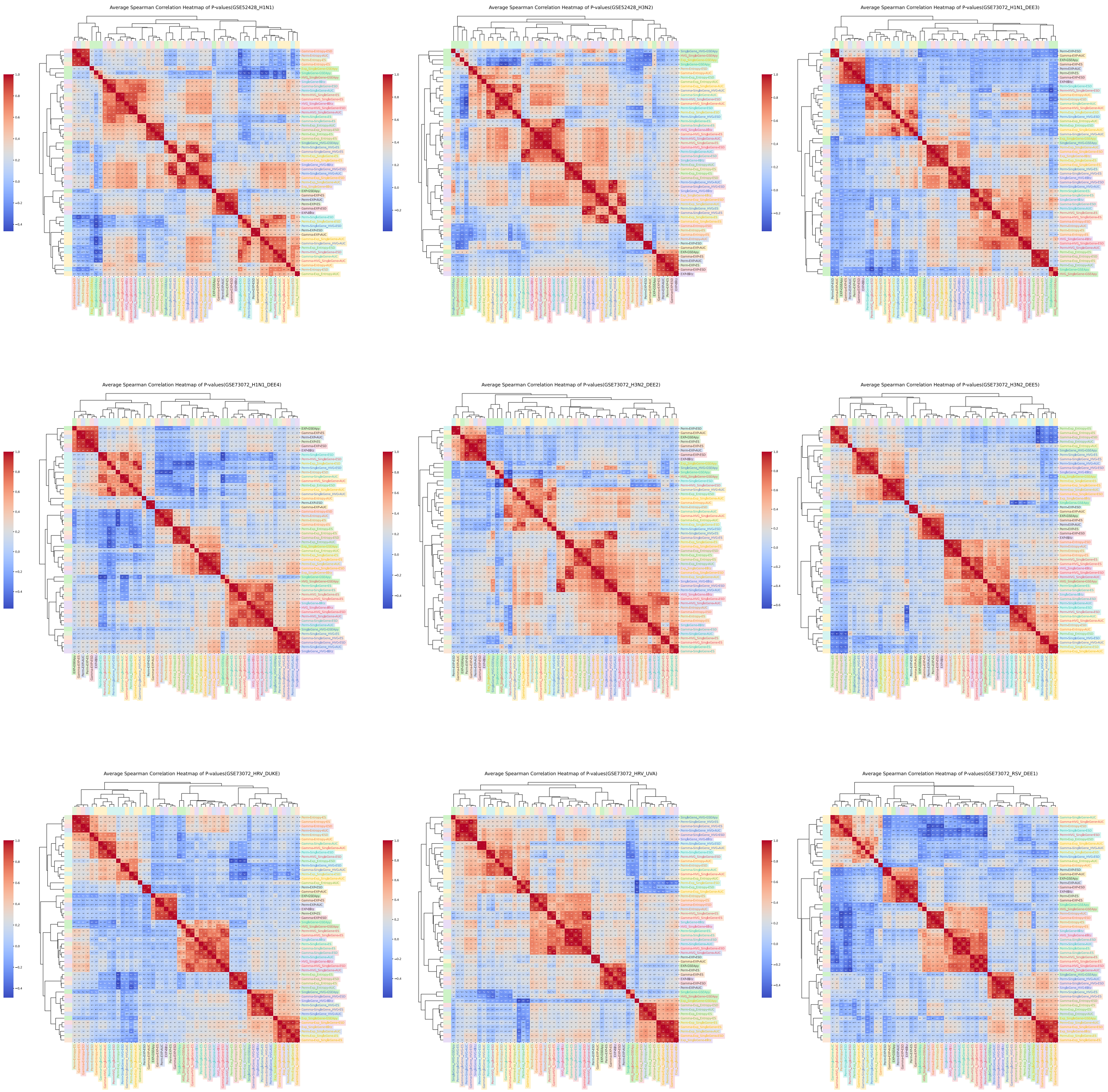

Supplementary Figure 3: Average Spearman correlation heatmap for different disease datasets. The supplementary heatmap provides a detailed breakdown visualization of the dataset presented in the main text across individual datasets, analyzing the similarity between each pair of methods under different permutations and repetitions. Each boxplot corresponds to a flu dataset within the combined dataset in the main text, including datasets GSE52428 with either influenza H1N1 or H3N2 viruses and GSE73072 with influenza A subtypes (H1N1 and H3N2), human rhinovirus (HRV), and respiratory syncytial virus (RSV).

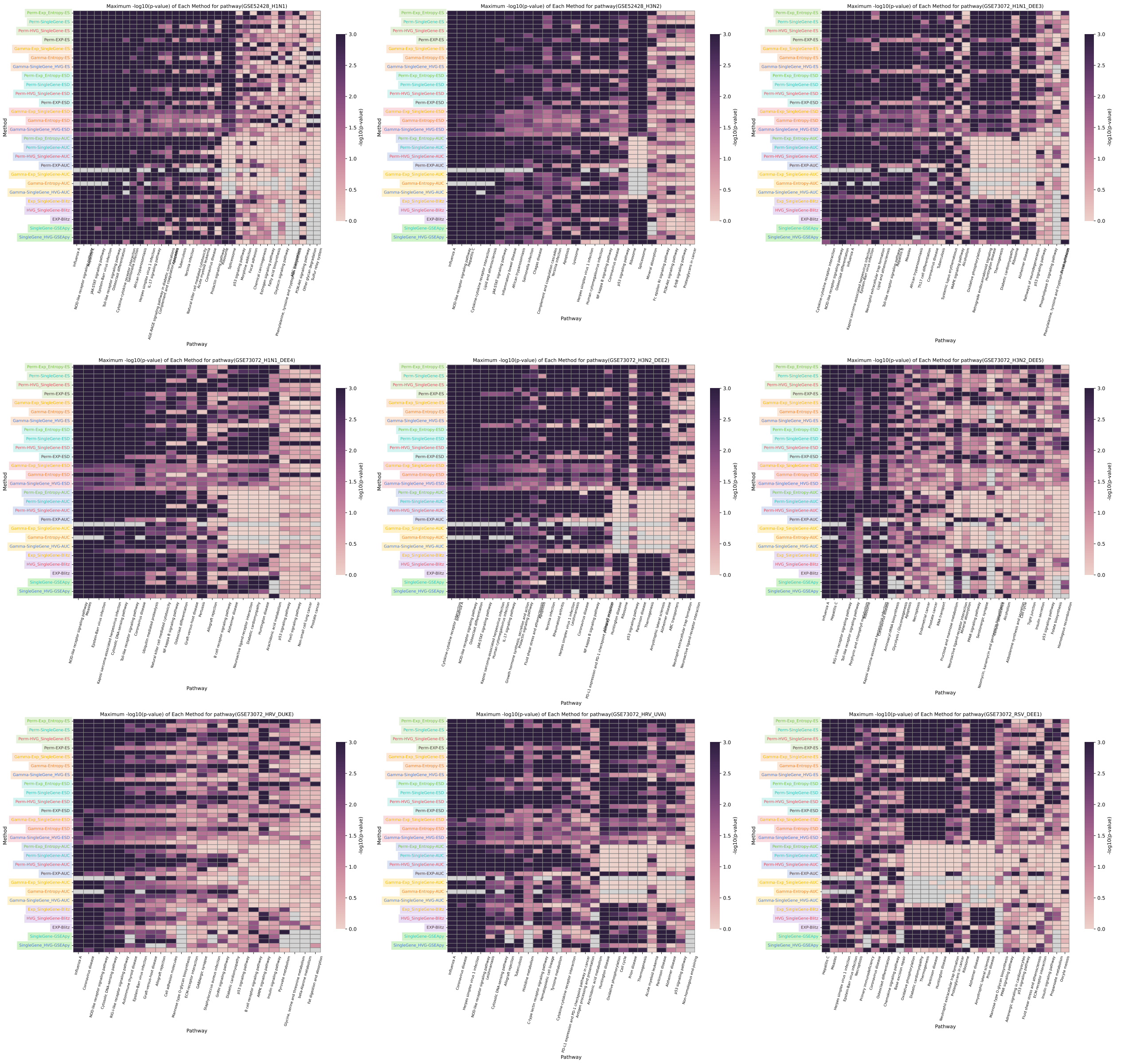

Supplementary Figure 4: Heatmap displaying the maximum $-\log_{10}(p\text{-value})$ obtained by each preprocessing method for its most significantly enriched pathway.

The supplementary heatmap displays the minimum $-\log_{10}(p\text{-value})$ obtained by each preprocessing method for its most significantly enriched pathway. The y-axis lists preprocessing methods, and the x-axis shows the corresponding pathways identified by each method. Color intensity reflects the statistical significance, with higher values indicating stronger enrichment. Each heatmap corresponds to a flu dataset within the combined dataset in the main text, including datasets GSE52428 with either influenza H1N1 or H3N2 viruses and GSE73072 with influenza A subtypes (H1N1 and H3N2), human rhinovirus (HRV), and respiratory syncytial virus (RSV).

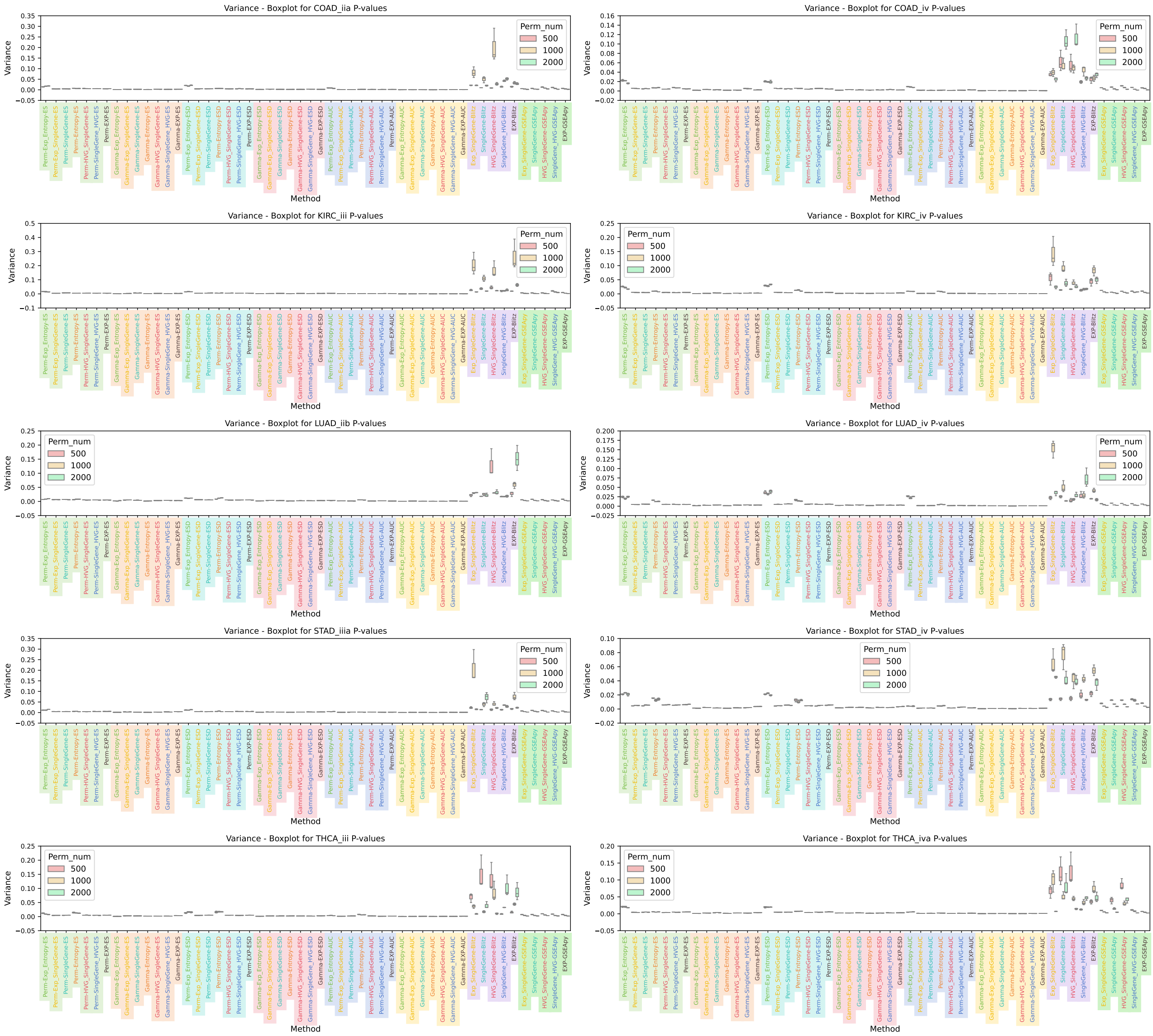

Supplementary Figure 5: Comparison of $p$-value variability among different approaches across three independent trials using 500, 1000, and 2000 permutations on various disease-related datasets.

The supplementary figure demonstrates the stability of $p$-value outputs under varying permutation counts across multiple cancer cohorts. Each boxplot reflects results from one stage of five tumor types evaluated in the study, namely colon (COAD), kidney (KIRC), lung (LUAD), stomach (STAD), and thyroid (THCA) carcinomas, providing a detailed comparison of methodological reproducibility across heterogeneous biological contexts.

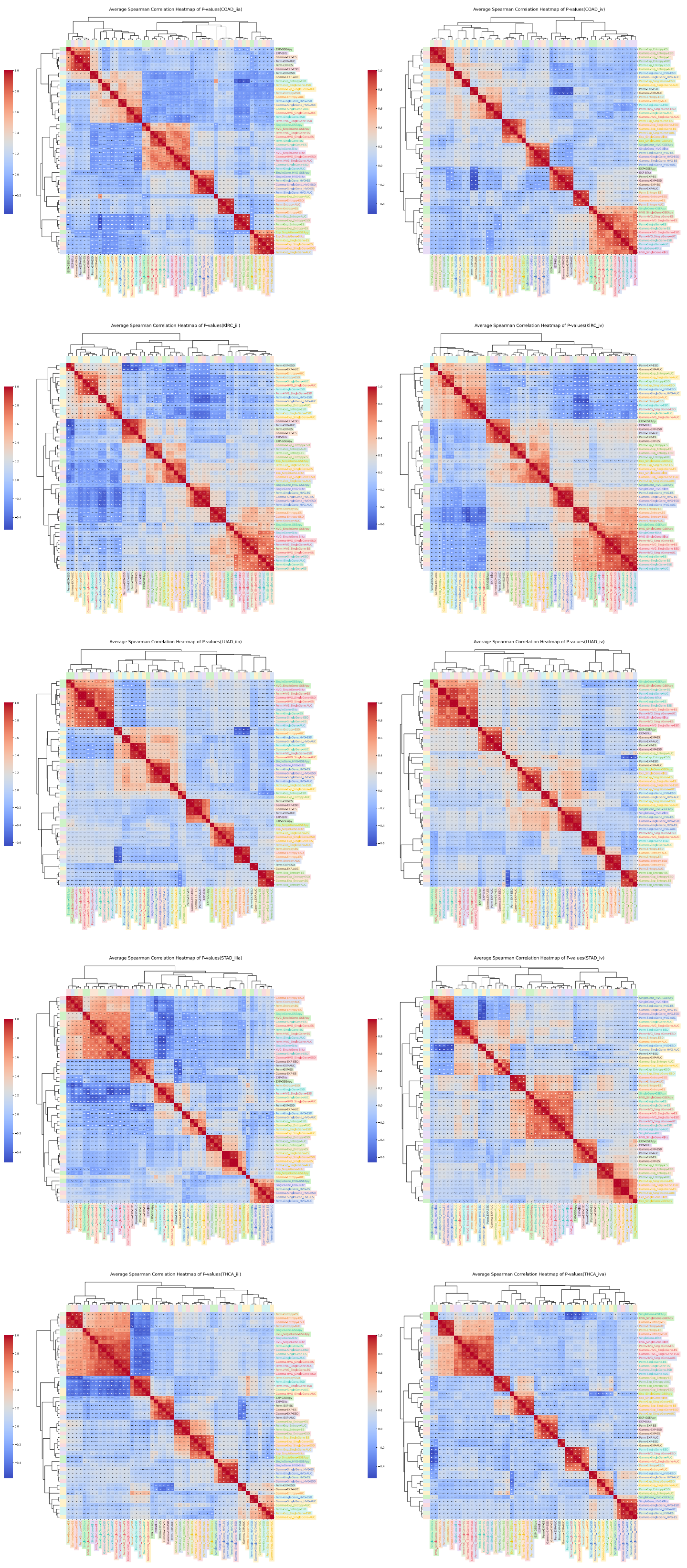

Supplementary Figure 6: Heatmap of Average Spearman Correlations Across Cancer Datasets
This heatmap summarizes the average pairwise correlations between methods across five cancer types—COAD, KIRC, LUAD, STAD, and THCA—under varying permutations and repetitions.

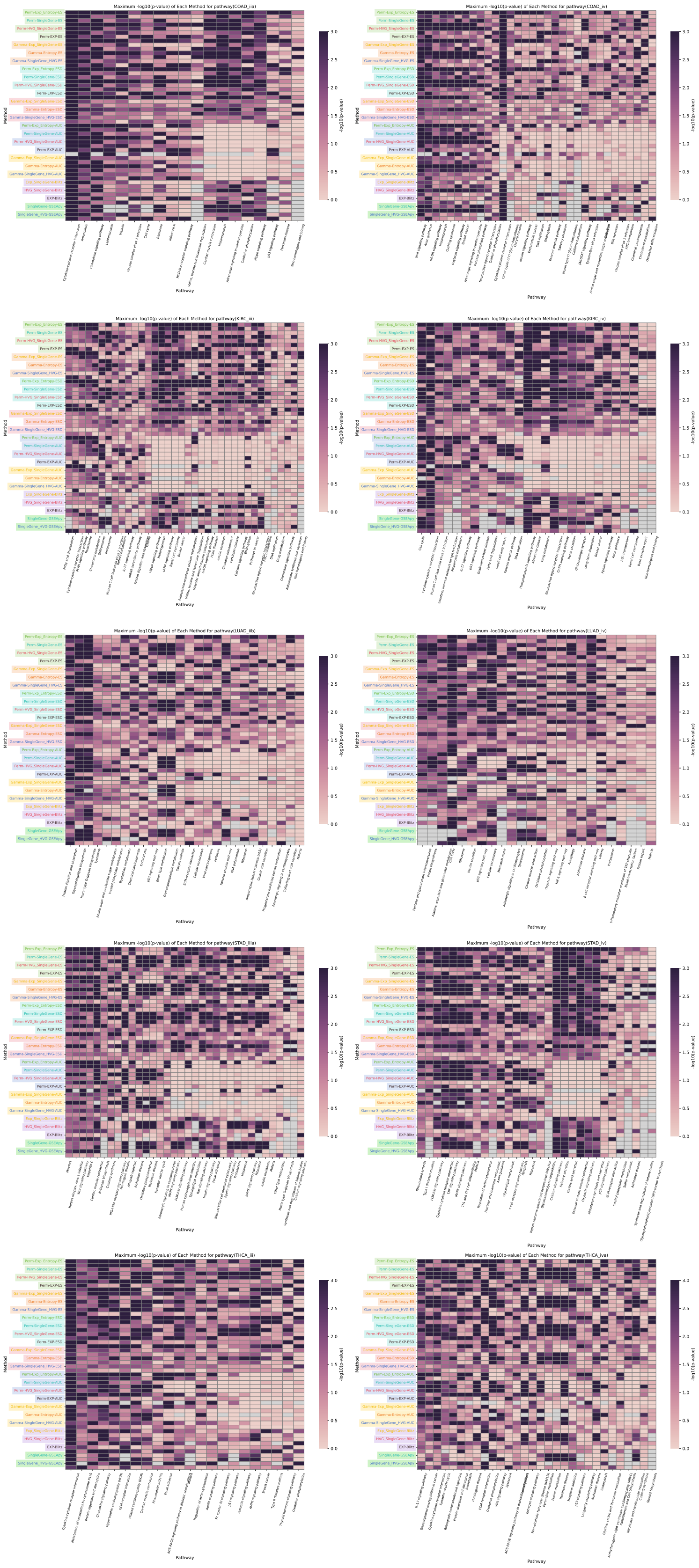

Supplementary Figure 7: Heatmap Showing the Peak $-\log_{10}(p\text{-value})$ for Top-Enriched Pathways Across Preprocessing Strategies
This heatmap presents the highest $-\log_{10}(p\text{-value})$ attained by each preprocessing method for its most significantly enriched pathway. The y-axis lists the preprocessing techniques, and the x-axis shows the corresponding top-ranked pathways. Color intensity denotes the level of statistical significance. The figure compares results across five cancer datasets—COAD, KIRC, LUAD, STAD, and THCA—revealing consistent performance advantages of our proposed method over existing approaches in terms of pathway detection stability.

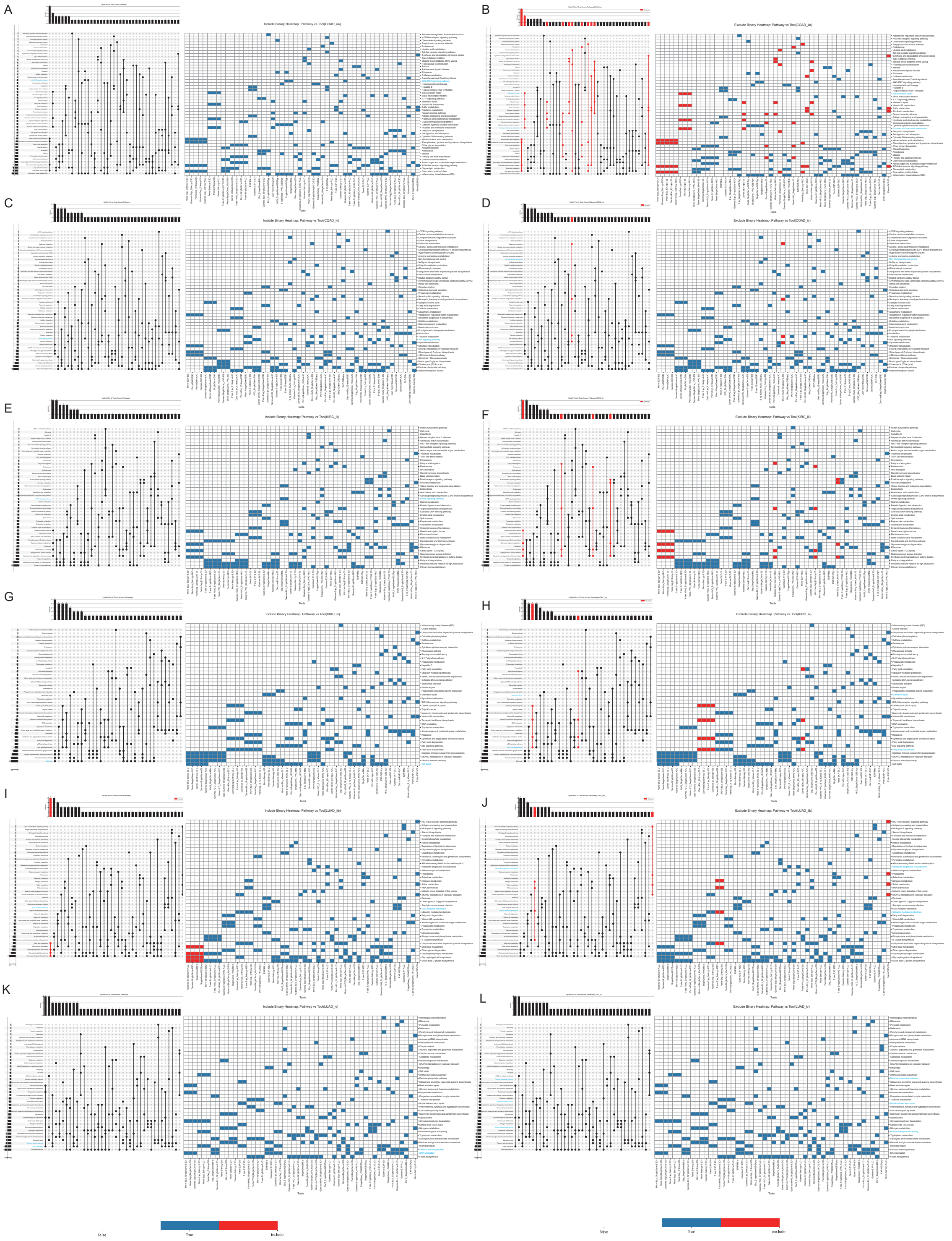

Supplementary Figure 8: (A, C, E, G, I, H) UpSet plots illustrate the overlap of the top 4 significantly enriched pathway combinations identified in COAD stage IIA, COAD stage IV, KIRC stage III, KIRC stage IV, LUAD stage IIA, and LUAD stage IV. The accompanying heatmaps display the specific methods associated with each pathway combination. Red highlights indicate combinations that include all three methods: GREA, blitzGSEA, and GSEApy. (B, D, F, H, J, L) UpSet plots show the top 4 significantly enriched pathway combination overlaps for COAD stage IIA, COAD stage IV, KIRC stage III, KIRC stage IV, LUAD stage IIA, and LUAD stage IV. In the corresponding heatmaps, red highlights mark pathway combinations that are identified solely by GREA, without contributions from blitzGSEA or GSEApy.

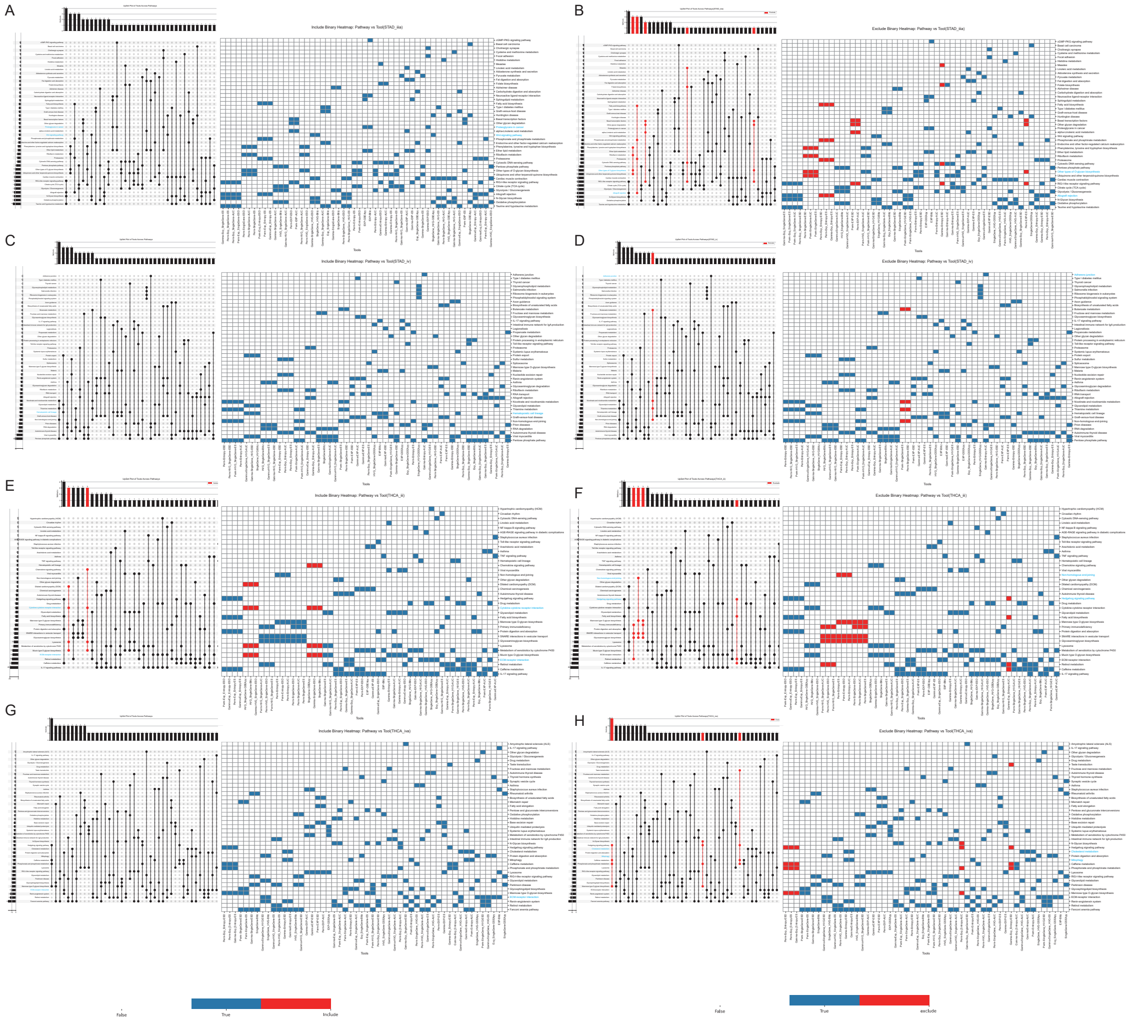

Supplementary Figure 9: (A, C, E, G) UpSet plots illustrate the overlap of the top 4 significantly enriched pathway combinations identified in STAD stage IIIA, STAD stage IV, THCA stage III, and THCA stage IVA. The accompanying heatmaps display the specific methods associated with each pathway combination. Red highlights indicate combinations that include all three methods: GREA, blitzGSEA, and GSEApy. (B, D, F, H) UpSet plots show the top 4 significantly enriched pathway combination overlaps for STAD stage IIIA, STAD stage IV, THCA stage III, and THCA stage IVA. In the corresponding heatmaps, red highlights mark pathway combinations that are identified solely by GREA, without contributions from blitzGSEA or GSEApy.

[14] Charles Spearman. The proof and measurement of association between two things. 1961.
